# Analysis of genes co-evolved with Igf2bp RNA-Binding Proteins provides insights into post-transcriptional regulatory networks

**DOI:** 10.64898/2026.03.05.709831

**Authors:** Joel K. Yisraeli, Sivan Izraely, Yuval Tabach

## Abstract

The Insulin-like growth factor 2 mRNA-binding protein (Igf2bp) family comprises three paralogous RNA-binding proteins—Igf2bp1, Igf2bp2, and Igf2bp3—that are highly conserved across chordates. Originally identified through diverse experimental screens probing intracellular RNA localization, RNA stability, and translational control, Igf2bps were initially studied in isolation from one another and within narrowly defined molecular contexts. Over time, it has become clear that these proteins act as multifunctional regulators of RNA metabolism and participate in a broad range of developmental and pathological processes. Here, we review the discovery, molecular functions, and biological roles of Igf2bp proteins, with particular emphasis on their conserved involvement in nervous system development and their reactivation in cancer. Given the emerging appreciation of the connection between genes involved in neural development and tumorigenesis, we thought it might be informative to perform a comparative evolutionary analysis to identify genes that coevolved with Igf2bp paralogs. These coevolved genes are strongly enriched for functions in axon guidance and neurogenesis, but also substantially overlap with experimentally defined Igf2bp1 RNA targets in lung adenocarcinoma (LUAD) cells. Notably, many of these genes are downregulated upon pharmacological inhibition of Igf2bp1 RNA binding. In this speculative review, we propose a model in which Igf2bp proteins evolved as coordinators of post-transcriptional gene regulation in the developing nervous system and were later co-opted in cancer to stabilize and coordinate oncogenic gene expression programs.

## Introduction

RNA-binding proteins (RBPs) are central regulators of post-transcriptional gene expression, influencing RNA localization, stability, translation, and processing. Among these, the Igf2bp family represents a striking example of how a single group of RBPs can integrate multiple modes of RNA regulation to shape cellular behavior. The family consists of three paralogs—Igf2bp1, Igf2bp2, and Igf2bp3—that share conserved domain architecture and are expressed dynamically during development and disease.

The discovery of Igf2bp proteins occurred through a series of independent experimental approaches, each highlighting a distinct molecular function. Igf2bp1 was first identified in chick embryo fibroblasts as Zipcode Binding Protein 1 (ZBP-1), required for the localization of β-actin mRNA to the leading edge of migrating cells (Ross et al., 1997). Igf2bp3 was cloned as a KH-domain-containing protein overexpressed in human pancreatic carcinoma (KOC) (Mueller-Pillasch et al., 1997). In *Xenopus*, the homolog of Igf2bp3 was isolated from oocytes as a factor binding the cis-acting element responsible for vegetal localization of Vg1 mRNA (Vera/Vg1RBP) (Deshler et al., 1998; Havin et al., 1998). Independently, the mammalian homolog of Igf2bp1 was identified as CRD-BP, a protein binding the coding region instability determinant of c-myc mRNA in a human erythroleukemia cell line (Doyle et al., 1998). Igf2bp2 was discovered using autoantibodies from hepatocellular carcinoma patients (p62) (Zhang et al., 1999). Finally, all three paralogs were cloned using the 5′ untranslated region of IGF-II leader 3 mRNA as bait in human rhabdomyosarcoma cells (IMP1–3) (Nielsen et al., 1999).

Much like the parable of the blind men and the elephant, these early studies each captured a different facet of Igf2bp function without revealing the full scope of their biological roles. Subsequent work has expanded the repertoire of Igf2bp-mediated regulation to include additional processes such as alternative splicing and recognition of m^6^A-modified transcripts (Degrauwe et al., 2016; Duan et al., 2024; Huang et al., 2018a). This multiplicity of mechanisms positions Igf2bps as integrators of post-transcriptional regulation rather than single-function RBPs.

### Molecular Functions of Igf2bp Proteins

Igf2bp proteins bind large and diverse sets of RNA targets, often in a context-dependent manner (Huang et al., 2018b; Muller et al., 2020). They can regulate RNA localization, as demonstrated by β-actin and Vg1 mRNAs (Deshler et al., 1998; Havin et al., 1998; Ross et al., 1997), stabilize transcripts such as c-myc (Doyle et al., 1998; Weidensdorfer et al., 2009), and modulate translation efficiency (Huttelmaier et al., 2005; Nielsen et al., 1999). More recently, Igf2bps have been implicated in reading m^6^A modifications and in influencing alternative splicing decisions (Degrauwe et al., 2016; Duan et al., 2024; Huang et al., 2018a). These activities are not mutually exclusive; rather, they allow Igf2bps to coordinate multiple aspects of RNA fate simultaneously.

A dominant feature of Igf2bp proteins is a capacity to regulate batteries of RNAs that participate in related cellular processes (Bell et al., 2013; Degrauwe et al., 2016; Wallis et al., 2025). This property distinguishes them from RBPs that act on limited sets of targets and suggests a systems-level role in shaping cellular phenotypes. In this regard it is possible to look on these proteins much as DNA transcription factors regulate batteries of target genes.

### Igf2bp proteins in development

The Igf2bp family of RBPs plays conserved and multifaceted roles during embryonic development, functioning primarily as post-transcriptional regulators that control RNA localization, stability, and translational timing. A unifying feature of Igf2bp proteins across developmental contexts is their repeated involvement in highly polarized cells and tissues, where spatially restricted protein synthesis is required to drive patterning, morphogenesis, and differentiation.

### Oogenesis and maternal RNA programs

The earliest and most mechanistically resolved developmental roles of Igf2bp proteins are found in oogenesis, where maternal transcripts must be selectively transported, anchored, and translationally regulated to establish embryonic axes and developmental competence. In *Xenopus*, Igf2bp3 was identified as a direct binding protein for the vegetal localization element of *Vg1* mRNA, a TGF-β family transcript essential for mesoderm induction and axis formation. Igf2bp3 associates with *Vg1* mRNA in oocytes and is required for its directed transport and stable localization at the vegetal cortex (Deshler et al., 1997; Havin et al., 1998). These findings build on earlier work demonstrating that *Vg1* localization depends on coordinated microtubule- and microfilament-based transport and anchoring pathways, defining the cellular framework within which Igf2bp3 operates (Yisraeli et al., 1990).

In *Drosophila*, the Igf2bp homolog Imp is a component of maternal ribonucleoprotein complexes and contributes to post-transcriptional regulation during oogenesis. Imp binds *oskar* mRNA and participates in its translational repression and anchoring, thereby helping to ensure proper germ plasm assembly and posterior patterning. Although Imp is not the sole determinant of *oskar* localization, it functions together with other RBPs to couple RNA localization with translational control during oocyte maturation (Munro et al., 2006). Together, these studies establish oogenesis as a foundational developmental context in which Igf2bp family proteins coordinate RNA localization and translational timing.

### Nervous system development

A second major developmental arena in which Igf2bp proteins play prominent roles is the nervous system, where neuronal differentiation, migration, and circuit assembly depend critically on localized RNA regulation. In vertebrates, Igf2bp1 is a canonical regulator of β-actin mRNA localization and translation, providing a mechanistic basis for polarized cytoskeletal remodeling and directed cell migration during development (Huttelmaier et al., 2005; Ross et al., 1997). In the developing nervous system, Igf2bp proteins appear to play both a cell autonomous and non-cell autonomous role in regulating cranial neural crest migration and epithelial integrity (Carmel et al., 2015; Yaniv et al., 2003). In developing neurons, Igf2bp1 localizes to axons and dendrites, associates with RNA granules, and regulates the transport and localized translation of mRNAs encoding cytoskeletal and signaling proteins. Through these activities, Igf2bp1 contributes to axon outgrowth, growth cone dynamics, and axon guidance, enabling neurons to respond rapidly and spatially to extracellular cues (Welshhans and Bassell, 2011). In hippocampal neurons, Igf2bp1 binds β-actin mRNA and undergoes activity-dependent trafficking within dendrites and dendritic spines; perturbing Igf2bp1 reduces dendritic β-actin mRNA localization and disrupts dendritic filopodia/synapse development (Eom et al., 2003; Perycz et al., 2011; Tiruchinapalli et al., 2003). Igf2bp1 knockout mice show disorganization in the developing neocortex along with decreased transport and anchoring of β-actin mRNA in neurons (Nunez et al., 2022). In the Xenopus optic tectum, Igf2bp3 is required for terminal arborization and promotes synaptic complexity by regulating mRNA translation (Kalous et al., 2014).

Developmental studies in *Drosophila* further underscore the evolutionary conservation of Igf2bp family function in the nervous system. The Igf2bp homolog Imp (Adolph et al., 2009) regulates neural stem cell growth and temporal patterning by stabilizing *myc* mRNA, thereby influencing neuroblast size and division rates during larval brain development (Samuels et al., 2020). Imp acts in opposition to the conserved RBP Syp/SYNCRIP as part of an intrinsic temporal program that governs neural stem cell decommissioning and the sequential generation of neuronal identities (Yang et al., 2017). In mushroom body neurons, Imp plays a critical role in developmental axon remodeling: it is transported into γ-Kenyon cell axons and is required for axonal regrowth and branching following pruning during metamorphosis, in part through post-transcriptional regulation of *chickadee/profilin* mRNA (Medioni et al., 2014). Most recently, in vivo RNA-interactome mapping demonstrated that Imp and Syp bind overlapping, temporally dynamic sets of mRNAs enriched for neurodevelopmental regulators, positioning Imp as a coordinator of time-dependent post-transcriptional networks during brain development (Lee et al., 2025). These findings complement earlier genetic work implicating Imp in neuronal migration and axon guidance and collectively establish Igf2bp family proteins as central regulators of nervous system development across species.

Taken together, developmental studies across oogenesis and neurogenesis support a coherent model in which Igf2bp proteins act as systems-level post-transcriptional organizers, coordinating the fate of batteries of RNAs in contexts requiring polarity, migration, and precise temporal control.

### Igf2bps in Cancer

There is an emerging appreciation of the role of neural developmental programs in cancer progression, demonstrating that tumors engage axon guidance cues, neurotrophic signaling, and neoneurogenesis mechanisms that mirror aspects of neurogenesis and neural circuit formation (Hanahan and Monje, 2023; Huang et al., 2025; Winkler et al., 2023). These processes can contribute directly to oncogenic growth, invasion, programming of the tumor microenvironment, and immune modulation. In addition, neurogenesis depends heavily on RNA localization and translational control, in part via RBPs that are particularly important in polarized cells. Many of the same RBPs that play a role in neural development are turned on or overexpressed in cancer cells, enabling coordination of entire gene expression programs post-transcriptionally.

In many different cancers, one or more of the Igf2bp proteins are upregulated (Bell et al., 2013; Duan et al., 2024; Nielsen et al., 1999). Across multiple cancer types, Igf2bps promote proliferation, suppress apoptosis, enhance migration and metastasis, confer chemoresistance, inhibit cell death, enable immune evasion, promote tumor metabolic reprogramming, and in general help maintain the oncogenic state (Bley et al., 2025; Degrauwe et al., 2016; Duan et al., 2024; Zhou et al., 2026). Although all three of the Igf2bp paralogs are expressed at various times and in various locations during embryogenesis, Igf2bp1 and Igf2bp3 expression is downregulated to nondetectable levels in virtually all tissues after birth, leading to the classification of Igf2bp1 and Igf2bp3 as oncofetal proteins.

A recurring theme is the context dependence of Igf2bp RNA targets. Different cancers display largely non-overlapping target sets, reflecting the cellular environment and available transcriptome (Huang et al., 2018b; Muller et al., 2020). This flexibility allows Igf2bps to be repurposed for distinct oncogenic programs while maintaining a conserved mode of action.

Several pharmacological inhibitors of Igf2bp1-RNA interactions have been reported over the last few years (Cai et al., 2024). BTYNB was identified in a fluorescence polarization screen as an inhibitor of Igf2bp1 binding to cMyc RNA and has been shown to inhibit several different types of cancers (Bley et al., 2025; Hagemann et al., 2023; Mahapatra et al., 2017; Schott et al., 2025; Xu et al., 2025). A11, a derivative of Cucurbitacin B, binds Igf2bp1 with high affinity, apparently in the KH1-KH2 domain, and inhibits A546 lung carcinoma cells in a xenograft model (Shang et al., 2023). AVJ16 (Singh et al., 2024) is an enhanced version of an Igf2bp1 inhibitor identified in a fluorescence polarization screen for inhibitors of Kras RNA binding (Wallis et al., 2022). AVJ16 destabilizes Igf2bp1-bound transcripts and suppresses tumor growth in colorectal carcinoma (CRC) and lung adenocarcinoma (LUAD) cell lines and in vivo in mouse xenografts (Singh et al., 2024; Wallis et al., 2025). These studies highlight the functional importance of Igf2bp-mediated RNA stabilization in cancer.

### Speculative hypothesis

The deployment of Igf2bp proteins in oocytes and neurons - two cell types that rely on RNA localization, stability, and translational regulation - suggests that these functions might represent ancestral roles of the family. These proteins are also activated in many neoplastic cells, where their functions are also of particular importance. If this is the case, then genes coevolving with Igf2bp proteins might represent a core network that could be regulated by the proteins as needed in different cellular settings. We find that the expression of Igf2bp1 coevolved genes is indeed strikingly enriched in neural developmental processes, including axonogenesis and axon guidance, and in different cancers and carcinogenesis. Using H1299 human LUAD cells as an initial proof of concept, we find that these coevolved genes encode RNAs that are enriched in Igf2bp1 binding and are also enriched in RNAs that are downregulated by pharmacological inhibition of Igf2bp1. These results support the hypothesis that the initial roles of Igf2bp proteins during development, and at least part of their core, regulated network, have been co-opted to regulate similar processes in cancer cells and their surroundings.

## Results

### Evolutionary Analysis of Igf2bp Coevolved Genes

To gain insights into the roles of Igf2bps and the targets with which they interact, we used the Cladeoscope algorithm to identify genes that coevolved with either Igf2bp1, 2, or 3. CladeOScope is a comparative genomics tool for the prediction of functional interactions between genes based on their phylogenetic profile, via a clade co-evolution prism. It analyzes the patterns of presence or absence of a particular gene with other genes across over a thousand different species to discover evolutionary relationships and functional associations. Identification of coevolving genes often reveals networks that function together and can highlight important interactions that can be leveraged for therapeutic purposes (Tsaban et al., 2021). It is important to note that the comparison is without regard to RNA transcription data. Figure 1 shows the 100 top genes that coevolved with Igf2bp1. Gene Set Enrichment Analysis revealed a strong and unexpected association between Igf2bp1 coevolved genes and genes expressed in pathways associated with nervous system development, particularly axon guidance and axonogenesis, as well as with different types of cancer (Figure 2A). As expected, given their high degree of conservation, the lists of genes that coevolved with Igf2bp2 and 3 were almost identical and yielded virtually the same list of enriched pathways as with Igf2bp1 (Supplemental Figure 1A, B). Multiple enriched gene sets converge on a shared subset of genes, indicating the presence of highly connected hubs within the coevolved gene network (Figure 2B). These hub genes are shared across several independent enrichment terms, suggesting that the coevolutionary signal is driven by a core group of genes involved in closely related biological processes rather than by unrelated functional modules. Because the coevolved gene networks for Igf2bp2 and 3 were also almost identical to that of Igf2bp1 (Supplemental Figure 1C, D), we continued our analysis with the Igf2bp1 coevolved genes (although the results were basically the same for the other coevolved genes as well; data not shown).

**Figure 1.**
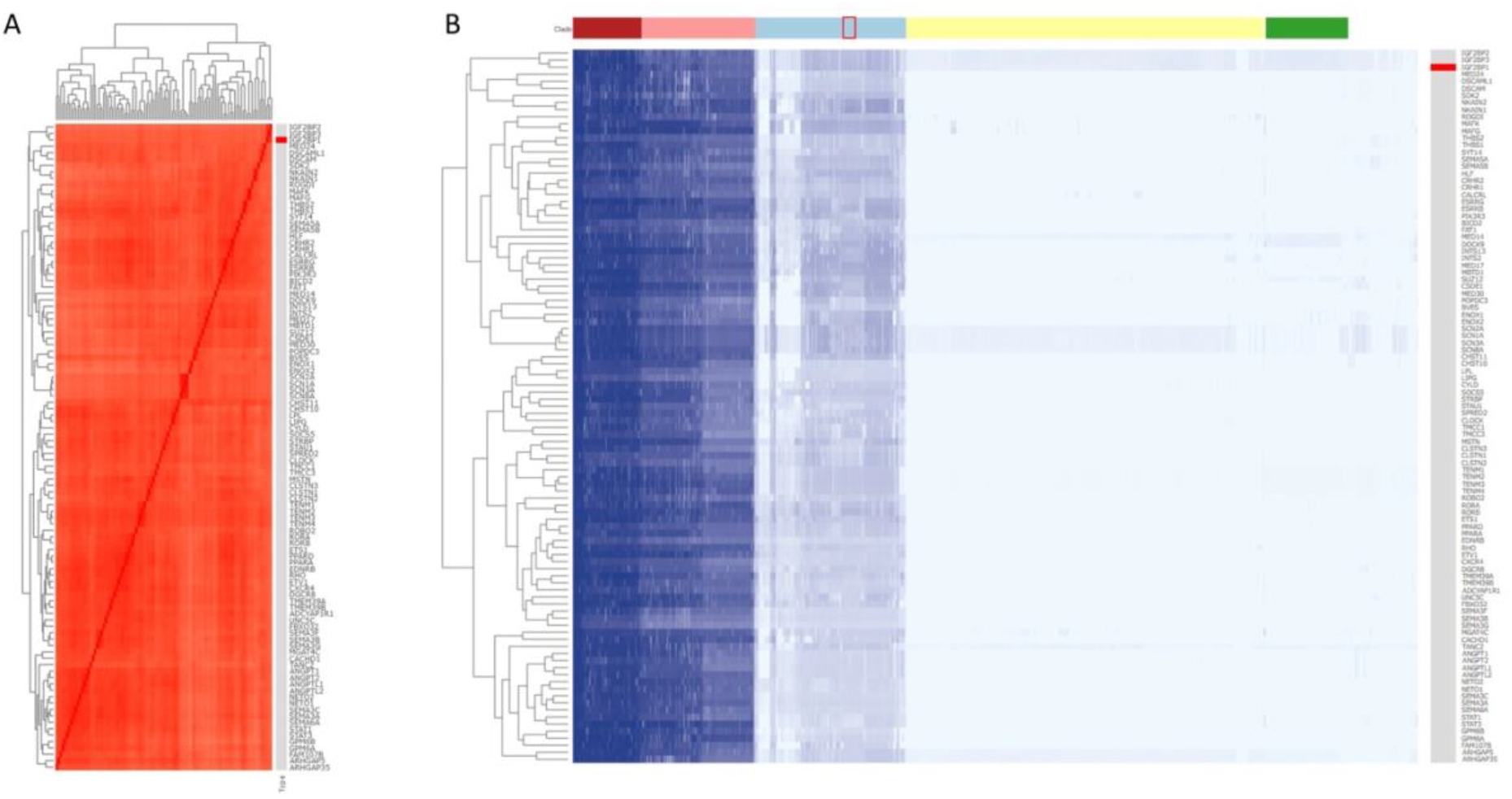
Genes that coevolved with Igf2bp1. (A) The heatmap, obtained by analysis with the CladeOScope algorithm (Tsaban et al., 2021), shows the correlations between the phylogenetic profiles of the output genes, given the Igf2bp1 gene as an input. The genes are clustered by their correlation patterns, as shown in the dendrogram at the top and left of the heatmap. Each pixel represents the cell’s value, the X-axis gene (column), and the Y axis-gene (row) comparison. (B) The heatmap, obtained by CladeOScope, shows the phylogenetic profiles of the output genes. The genes are clustered by their correlation patterns, as shown in the dendrogram on the leftmost of the heatmap. Each row corresponds to a gene, and each column to a species. Each pixel depicts the ‘Evolutionary Conservation’ score of gene Y in species X. Colors illustrate the conservation values; darker (blue) means a higher value. Each pixel represents the cell’s value, the Y-axis gene (row) and the X-axis species (column). The red rectangle shows the location of *Drosophila*.

**Figure 2.**
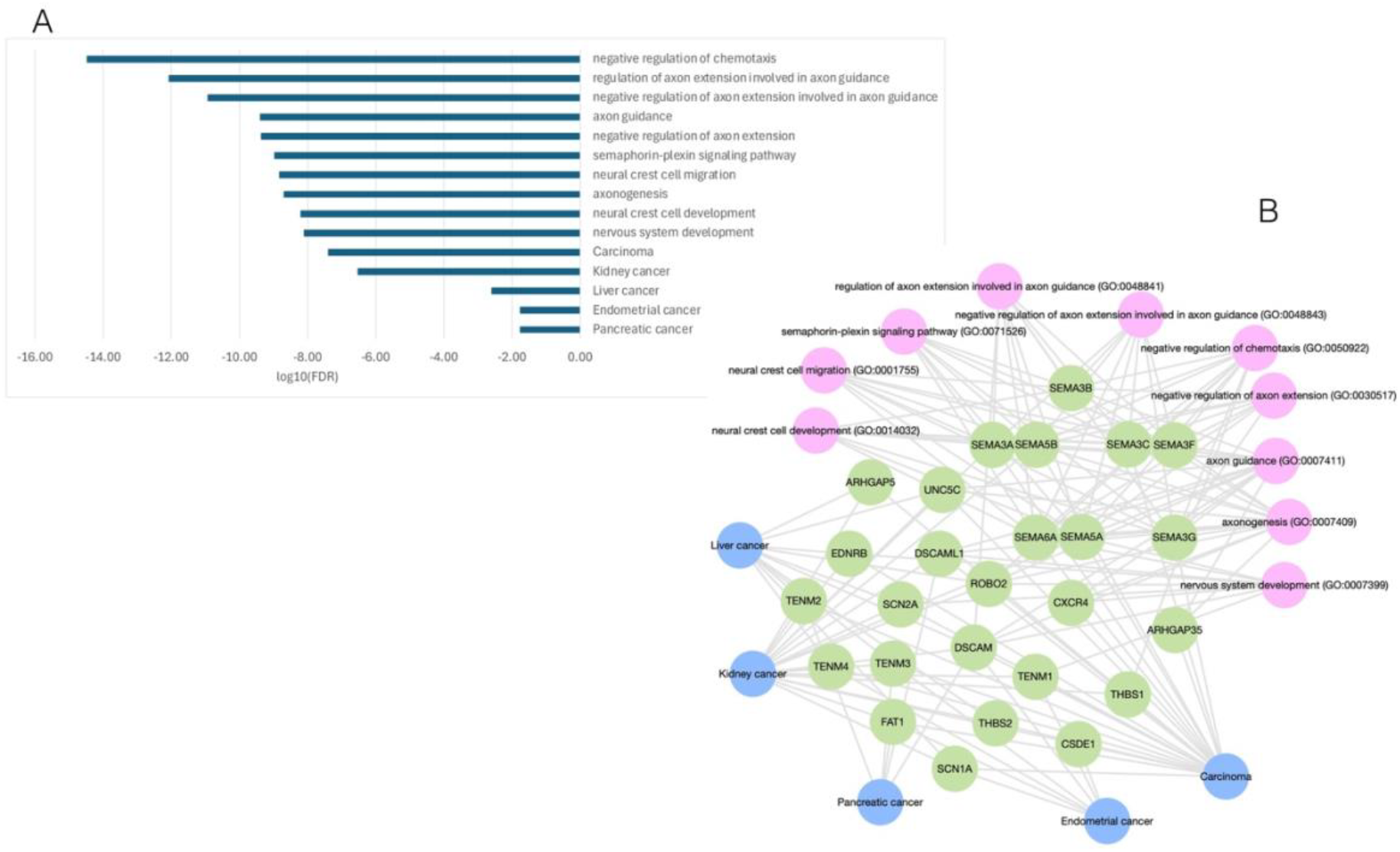
Gene set enrichment analysis of Igf2bp1 coevolved genes. (A) Gene set enrichment analysis was performed using EnrichR KG (Evangelista et al., 2023) on the 100 top Igf2bp1 coevolved genes, based on GO Biological Process and Jansen Disease libraries. (B) Network analysis (by the EnrichR KG program) of the 100 top coevolved genes shows the genes (hubs, in green) shared by multiple enriched gene sets (GO Biological Process, in pink; Jansen Diseases, in blue).

### Coevolved genes are enriched for direct Igf2bp1 RNA targets

To determine whether coevolved genes are also direct molecular targets of Igf2bp1, we compared the coevolved gene set to genes we previously identified as Igf2bp1-bound transcripts, in H1299 human LUAD cells, by using the eCLIP method to scan for RNAs crosslinked to Igf2bp1 (Wallis et al., 2025). Of the 100 genes identified as coevolved with *Igf2bp1*, 38 were independently identified as Igf2bp1 eCLIP targets (Figure 3A). This overlap represents a 1.61-fold enrichment over chance expectation and was statistically significant (one-tailed hypergeometric test, p = 8.4 × 10^−4^). These results indicate that the RNAs transcribed from genes coevolving with *Igf2bp1* are direct targets of Igf2bp1 in LUAD cells.

**Figure 3.**
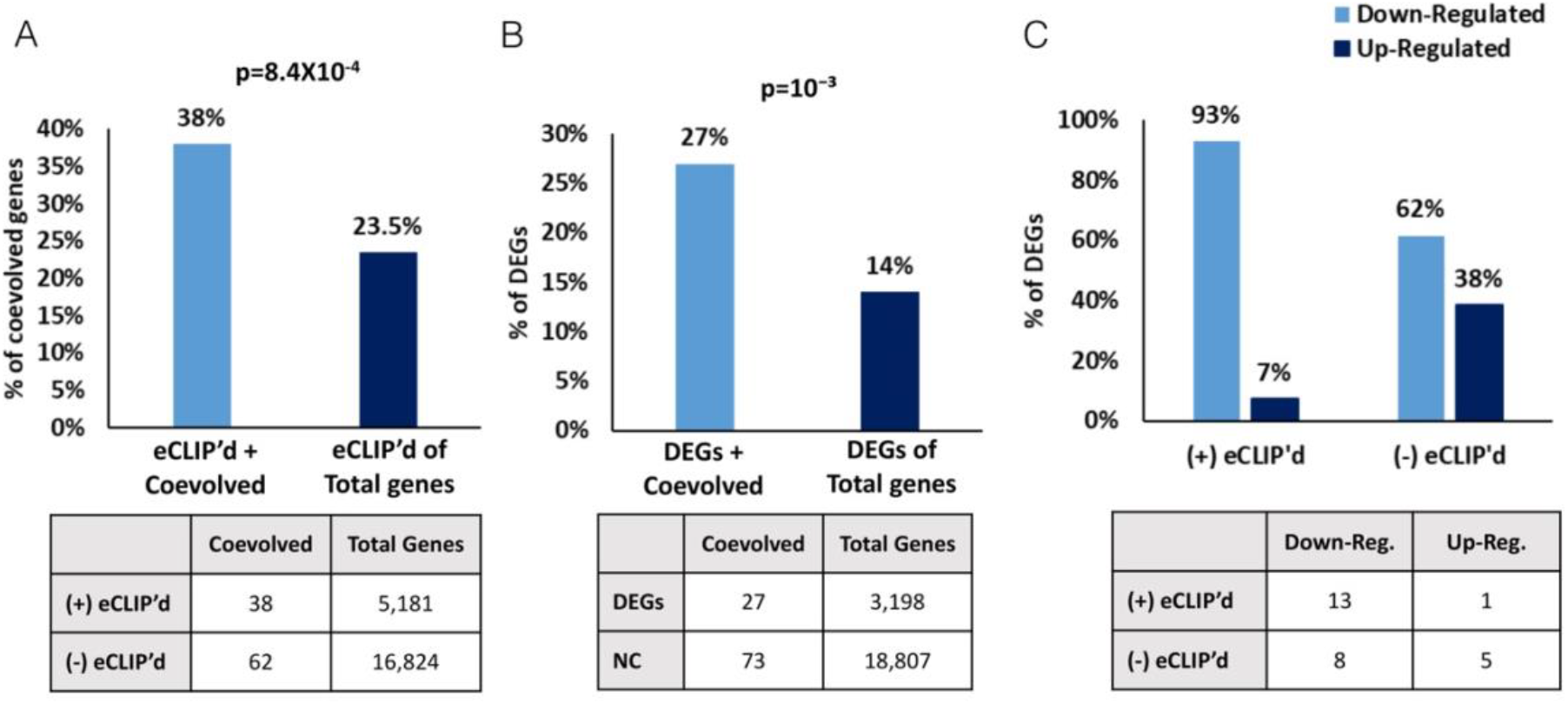
Correlation of coevolved genes with Igf2bp1 binding and pharmacological inhibition. (A) *Comparison of the Igf2bp1 coevolved gene set with the set of previously identified Igf2bp1-bound transcripts (based on an eCLIP analysis (Wallis et al., 2025)*. Of the 100 genes identified as coevolved with *Igf2bp1*, 38 were independently identified as Igf2bp1 eCLIP targets (2×2 contingency matrix, below). To test whether Igf2bp1-coevolved genes were enriched among Igf2bp1 eCLIP targets, we performed a one-tailed hypergeometric test using the set of RNA-seq-detected genes as the background universe (N = 22,005). The set of Igf2bp1 eCLIP targets comprised K = 5,181 genes, the coevolved set contained n = 100 genes, and the observed overlap was k = 38. The enrichment p-value was computed as P (X ≥ 38) for *X ∼* Hypergeometric (*N* = 22005, *K* = 5181, *n* = 100), yielding p = 8.4×10^−4^. The observed overlap represents a 1.61-fold enrichment over chance expectation (expected overlap 23.5 genes). (B) *Comparison of the Igf2bp1 coevolved gene set with the set of previously identified transcripts in H1299 cells that are responsive to AVJ16 (Wallis et al., 2025)*. Of the 100 coevolved genes, 27 were significantly responsive to AVJ16 treatment (2×2 contingency matrix, below). This represents an approximately 2.2-fold enrichment of drug-responsive genes relative to the transcriptome-wide response (one-tailed hypergeometric test, p ∼ 10^−3^). (C) *Analysis of the Igf2bp1 coevolved genes that are regulated by AVJ16*. Of the 27 coevolved genes whose expression was differentially regulated by AVJ16, 14 were direct targets of Igf2bp1, and 13 of these were downregulated (2×2 contingency matrix, below). Among the 13 non-eCLIP genes, only 8 were downregulated. This association was statistically significant (one-tailed Fisher’s exact test, p ≈ 0.045). DEG, differentially expressed genes; NC, not changed

### Coevolved Igf2bp1 targets are preferentially responsive to pharmacological inhibition

We next asked whether genes coevolved with *Igf2bp1* are functionally responsive to pharmacological disruption of Igf2bp1–RNA interactions. Treatment of H1299 cells with AVJ16, a small-molecule inhibitor of Igf2bp1 RNA binding that as we previously showed caused differential expression of a large number of RNAs (Wallis et al., 2025), resulted in differential expression of a substantial fraction of the coevolved gene set. Of the 100 coevolved genes, 27 were significantly responsive to AVJ16 treatment, with 21 exhibiting downregulation (Figure 3B). This represents an approximately 2.2-fold enrichment of drug-responsive genes relative to the transcriptome-wide response (p ∼ 10^−3^), consistent with destabilization of Igf2bp1-bound transcripts upon inhibition of RNA binding.

To further explore the connection between AVJ16-responsive RNAs and the coevolved genes, we compared those directly bound by Igf2bp1 (eCLIP-positive) as opposed to those not directly bound (non-eCLIPd RNAs). The eCLIPd RNAs were strongly biased toward downregulation following AVJ16 treatment (13/14; Figure 3C). Non-eCLIPd genes showed a more balanced response (8/13 downregulated; Figure 3C). This association was statistically significant (one-tailed Fisher’s exact test, p ≈ 0.045) and supports a model in which Igf2bp1 binding directly stabilizes a conserved subset of transcripts. Disruption of this interaction tends to lead to their selective degradation.

## Discussion

The convergence of evolutionary, functional, and pharmacological data presented here supports a unifying model for Igf2bp function. Using a comparative genomic approach, we identify a set of genes that coevolved with the Igf2bp paralogs and show that these genes are strongly enriched for pathways involved in nervous system development, particularly axon guidance and axonogenesis (Figure 2). The connection between Igf2bp and neurogenesis extends even beyond chordates to a distantly related Drosophila member of the family ((Lee et al., 2025; Medioni et al., 2014); Figure 1). Network analysis further reveals that this enrichment is driven by a core group of highly connected hub genes shared across multiple functional categories, suggesting coordinated evolutionary constraint rather than coincidental association.

Importantly, this evolutionary signal is reflected at the molecular level. A substantial fraction of the Igf2bp1-coevolved genes is independently identified as direct Igf2bp1 RNA targets in our eCLIP data, with the observed overlap significantly exceeding chance expectation (Figure 3A). This enrichment suggests that coevolution with Igf2bp1 is likely to be linked to direct RNA-protein interactions, supporting the idea that evolutionary pressure has acted on both the RBP and a subset of its RNA targets as a functional unit.

Pharmacological perturbation of Igf2bp1 RNA binding further connects these evolutionary and molecular observations to functional outcomes. Inhibition of Igf2bp1 by AVJ16 preferentially affects genes within the coevolved set, with a strong bias toward transcript downregulation (Figure 3B). Notably, this effect is most pronounced among genes that are both coevolved with Igf2bp1 and directly bound by Igf2bp1, the majority of which are destabilized upon AVJ16 treatment (Figure 3C). These findings are consistent with a model in which Igf2bp1 stabilizes a conserved subset of transcripts, and disruption of this interaction leads to their selective degradation.

Together, these results suggest that Igf2bp proteins may have originally evolved to coordinate post-transcriptional regulation of gene networks essential for nervous system development. The coevolution of Igf2bps with axon guidance-related genes, combined with their conserved expression and functional requirement in the developing nervous system, supports this interpretation. In cancer, this ancestral regulatory capacity appears to be redeployed to stabilize oncogenic gene expression programs, enabling tumor cells to exploit a deeply conserved post-transcriptional regulatory mechanism.

More broadly, these findings highlight how RBPs can function as systems-level regulators analogous to transcription factors, coordinating the expression of batteries of functionally related genes. The integration of evolutionary analysis with RNA-binding and pharmacological data provides a framework for identifying functionally critical RBP-RNA interactions and suggests that evolutionary constraint may serve as a useful predictor of therapeutic vulnerability in cancers driven by post-transcriptional dysregulation. The approach outlined here can function as a pipeline for analysing other families of RBPs, enabling discovery of hidden connections to developmental and disease processes.

We propose that Igf2bp proteins function as evolutionarily conserved post-transcriptional organizers, originally optimized to regulate complex, spatially organized gene expression programs in the developing nervous system. Their modular RNA-binding capabilities and responsiveness to cellular context make them particularly susceptible to pathological co-option in cancer. This model helps reconcile the apparent diversity of Igf2bp functions across development and disease and suggests that disruption of conserved regulatory Igf2bp-RNA interactions could provide novel directions for therapeutic interventions.

## Figures

**Supplemental Figure 1.**
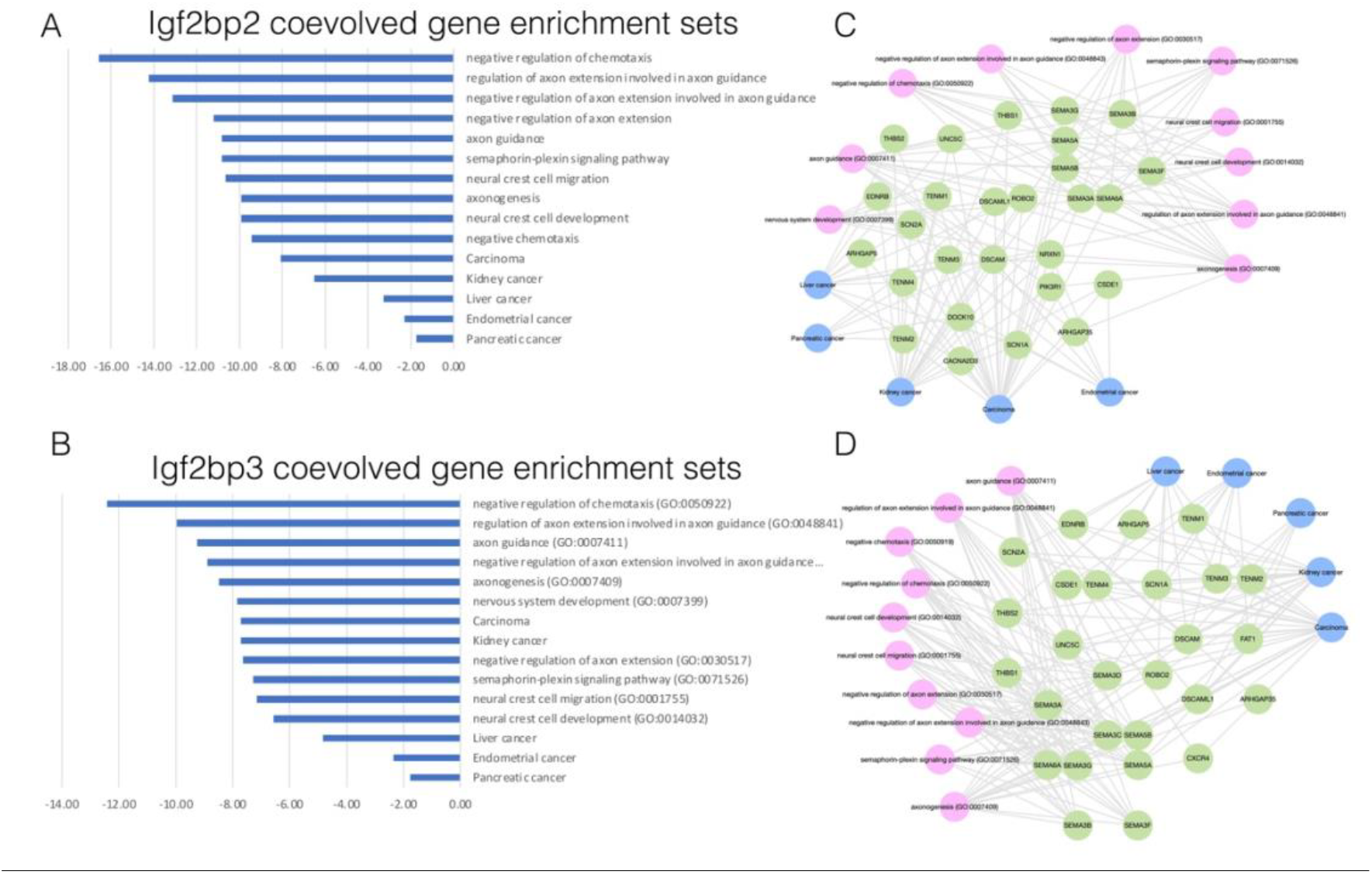
Gene set enrichment analysis of Igf2bp2 and Igf2bp3 coevolved genes. Gene set enrichment analysis was performed using EnrichR KG (Evangelista et al., 2023) on the 100 top Igf2bp2 (A) and Igf2bp3 (B) coevolved genes (obtained using the CladeOScope algorithm (Tsaban et al., 2021)), based on GO Biological Process and Jansen Disease libraries. Network analysis (by the EnrichR KG program) of the 100 top genes coevolved with Igf2bp2 (C) and Igf2bp3 (D) shows the genes (hubs, in green) shared by multiple enriched gene sets (GO Biological Process, in pink; Jansen Diseases, in blue).

## Acknowledgements

I would like to thank Prof. Hannah Margalit for helpful discussions and Dr. Froma Oberman for helpful comments on the manuscript. The work was supported by grants from the Israel Science Foundation and The Lung Cancer Research Foundation (LCRF) in partnership with the Israel Cancer Research Fund (JKY). JKY is the recipient of the Morley Goldblatt Chair in Cancer Research and Experimental Medicine.

## References

Adolph, S.K., DeLotto, R., Nielsen, F.C., and Christiansen, J. (2009). Embryonic expression of Drosophila IMP in the developing CNS and PNS. Gene Expr Patterns 9, 138–143.

Bell, J.L., Wachter, K., Muhleck, B., Pazaitis, N., Kohn, M., Lederer, M., and Huttelmaier, S. (2013). Insulin-like growth factor 2 mRNA-binding proteins (IGF2BPs): post-transcriptional drivers of cancer progression? Cell Mol Life Sci 70, 2657–2675.

Bley, N., Rausch, A., Muller, S., Simon, T., Glass, M., Misiak, D., Schian, L., Peters, L.M., Dipto, M., Hmedat, A., et al. (2025). Inhibition of RNA-binding proteins enhances immunotherapy in ovarian cancer. Signal Transduct Target Ther 10, 419.

Cai, Y., Wang, Y., Mao, B., You, Q., and Guo, X. (2024). Targeting insulin-like growth factor 2 mRNA-binding proteins (IGF2BPs) for the treatment of cancer. Eur J Med Chem 268, 116241.

Carmel, M.S., Kahane, N., Oberman, F., Miloslavski, R., Sela-Donenfeld, D., Kalcheim, C., and Yisraeli, J.K. (2015). A Novel Role for VICKZ Proteins in Maintaining Epithelial Integrity during Embryogenesis. PLoS One 10, e0136408.

Degrauwe, N., Suva, M.L., Janiszewska, M., Riggi, N., and Stamenkovic, I. (2016). IMPs: an RNA-binding protein family that provides a link between stem cell maintenance in normal development and cancer. Genes Dev 30, 2459–2474.

Deshler, J.O., Highett, M.I., Abramson, T., and Schnapp, B.J. (1998). A highly conserved RNA-binding protein for cytoplasmic mRNA localization in vertebrates. Curr Biol 8, 489–496.

Deshler, J.O., Highett, M.I., and Schnapp, B.J. (1997). Localization of Xenopus Vg1 mRNA by Vera protein and the endoplasmic reticulum. Science 276, 1128–1131.

Doyle, G.A., Betz, N.A., Leeds, P.F., Fleisig, A.J., Prokipcak, R.D., and Ross, J. (1998). The c-myc coding region determinant-binding protein: a member of a family of KH domain RNA-binding proteins. Nucleic Acids Res 26, 5036–5044.

Duan, M., Liu, H., Xu, S., Yang, Z., Zhang, F., Wang, G., Wang, Y., Zhao, S., and Jiang, X. (2024). IGF2BPs as novel m(6)A readers: Diverse roles in regulating cancer cell biological functions, hypoxia adaptation, metabolism, and immunosuppressive tumor microenvironment. Genes Dis 11, 890–920.

Eom, T., Antar, L.N., Singer, R.H., and Bassell, G.J. (2003). Localization of a beta-actin messenger zipcode-binding protein modulates the density of filopodial synapses. J Neurosci 23, 10433–10444.

Evangelista, J.E., Xie, Z., Marino, G.B., Nguyen, N., Clarke, D.J.B., and Ma’ayan, A. (2023). Enrichr-KG: bridging enrichment analysis across multiple libraries. Nucleic Acids Res 51, W168–W179.

Hagemann, S., Misiak, D., Bell, J.L., Fuchs, T., Lederer, M.I., Bley, N., Hammerle, M., Ghazy, E., Sippl, W., Schulte, J.H., et al. (2023). IGF2BP1 induces neuroblastoma via a druggable feedforward loop with MYCN promoting 17q oncogene expression. Mol Cancer 22, 88.

Hanahan, D., and Monje, M. (2023). Cancer hallmarks intersect with neuroscience in the tumor microenvironment. Cancer Cell 41, 573–580.

Havin, L., Git, A., Elisha, Z., Oberman, F., Yaniv, K., Schwartz, S.P., Standart, N., and Yisraeli, J.K. (1998). RNA-binding protein conserved in both microtubule- and microfilament-based RNA localization. Genes Dev 12, 1593–1598.

Huang, H., Weng, H., Sun, W., Qin, X., Shi, H., Wu, H., Zhao, B.S., Mesquita, A., Liu, C., Yuan, C.L., et al. (2018a). Recognition of RNA N(6)-methyladenosine by IGF2BP proteins enhances mRNA stability and translation. Nat Cell Biol 20, 285–295.

Huang, Q., Hu, B., Zhang, P., Yuan, Y., Yue, S., Chen, X., Liang, J., Tang, Z., and Zhang, B. (2025). Neuroscience of cancer: unraveling the complex interplay between the nervous system, the tumor and the tumor immune microenvironment. Mol Cancer 24, 24.

Huang, X., Zhang, H., Guo, X., Zhu, Z., Cai, H., and Kong, X. (2018b). Insulin-like growth factor 2 mRNA-binding protein 1 (IGF2BP1) in cancer. J Hematol Oncol 11, 88.

Huttelmaier, S., Zenklusen, D., Lederer, M., Dictenberg, J., Lorenz, M., Meng, X., Bassell, G.J., Condeelis, J., and Singer, R.H. (2005). Spatial regulation of beta-actin translation by Src-dependent phosphorylation of ZBP1. Nature 438, 512–515.

Kalous, A., Stake, J.I., Yisraeli, J.K., and Holt, C.E. (2014). RNA-binding protein Vg1RBP regulates terminal arbor formation but not long-range axon navigation in the developing visual system. Dev Neurobiol 74, 303–318.

Lee, J.Y., Huang, N., Samuels, T.J., and Davis, I. (2025). Imp/IGF2BP and Syp/SYNCRIP temporal RNA interactomes uncover combinatorial networks of regulators of Drosophila brain development. Sci Adv 11, eadr6682.

Mahapatra, L., Andruska, N., Mao, C., Le, J., and Shapiro, D.J. (2017). A Novel IMP1 Inhibitor, BTYNB, Targets c-Myc and Inhibits Melanoma and Ovarian Cancer Cell Proliferation. Transl Oncol 10, 818–827.

Medioni, C., Ramialison, M., Ephrussi, A., and Besse, F. (2014). Imp promotes axonal remodeling by regulating profilin mRNA during brain development. Curr Biol 24, 793–800.

Mueller-Pillasch, F., Lacher, U., Wallrapp, C., Micha, A., Zimmerhackl, F., Hameister, H., Varga, G., Friess, H., Buchler, M., Beger, H.G., et al. (1997). Cloning of a gene highly overexpressed in cancer coding for a novel KH-domain containing protein. Oncogene 14, 2729–2733.

Muller, S., Bley, N., Busch, B., Glass, M., Lederer, M., Misiak, C., Fuchs, T., Wedler, A., Haase, J., Bertoldo, J.B., et al. (2020). The oncofetal RNA-binding protein IGF2BP1 is a druggable, post-transcriptional super-enhancer of E2F-driven gene expression in cancer. Nucleic Acids Res 48, 8576–8590.

Munro, T.P., Kwon, S., Schnapp, B.J., and St Johnston, D. (2006). A repeated IMP-binding motif controls oskar mRNA translation and anchoring independently of Drosophila melanogaster IMP. J Cell Biol 172, 577–588.

Nielsen, J., Christiansen, J., Lykke-Andersen, J., Johnsen, A.H., Wewer, U.M., and Nielsen, F.C. (1999). A family of insulin-like growth factor II mRNA-binding proteins represses translation in late development. Mol Cell Biol 19, 1262–1270.

Nunez, L., Buxbaum, A.R., Katz, Z.B., Lopez-Jones, M., Nwokafor, C., Czaplinski, K., Pan, F., Rosenberg, J., Monday, H.R., and Singer, R.H. (2022). Tagged actin mRNA dysregulation in IGF2BP1-/-mice. Proc Natl Acad Sci U S A 119, e2208465119.

Perycz, M., Urbanska, A.S., Krawczyk, P.S., Parobczak, K., and Jaworski, J. (2011). Zipcode binding protein 1 regulates the development of dendritic arbors in hippocampal neurons. J Neurosci 31, 5271–5285.

Ross, A.F., Oleynikov, Y., Kislauskis, E.H., Taneja, K.L., and Singer, R.H. (1997). Characterization of a beta-actin mRNA zipcode-binding protein. Mol Cell Biol 17, 2158–2165.

Samuels, T.J., Jarvelin, A.I., Ish-Horowicz, D., and Davis, I. (2020). Imp/IGF2BP levels modulate individual neural stem cell growth and division through myc mRNA stability. Elife 9.

Schott, A., Simon, T., Muller, S., Rausch, A., Busch, B., Glass, M., Misiak, D., Dipto, M., Elrewany, H., Peters, L.M., et al. (2025). The IGF2BP1 oncogene is a druggable m(6)A-dependent enhancer of YAP1-driven gene expression in ovarian cancer. NAR Cancer 7, zcaf006.

Shang, F.F., Lu, Q., Lin, T., Pu, M., Xiao, R., Liu, W., Deng, H., Guo, H., Quan, Z.S., Ding, C., et al. (2023). Discovery of Triazolyl Derivatives of Cucurbitacin B Targeting IGF2BP1 against Non-Small Cell Lung Cancer. J Med Chem 66, 12931–12949.

Singh, A., Singh, V., Wallis, N., Abis, G., Oberman, F., Wood, T., Dhamdhere, M., Gershon, T., Ramos, A., Yisraeli, J., et al. (2024). Development of a specific and potent IGF2BP1 inhibitor: A promising therapeutic agent for IGF2BP1-expressing cancers. Eur J Med Chem 263, 115940.

Tiruchinapalli, D.M., Oleynikov, Y., Kelic, S., Shenoy, S.M., Hartley, A., Stanton, P.K., Singer, R.H., and Bassell, G.J. (2003). Activity-dependent trafficking and dynamic localization of zipcode binding protein 1 and beta-actin mRNA in dendrites and spines of hippocampal neurons. J Neurosci 23, 3251–3261.

Tsaban, T., Stupp, D., Sherill-Rofe, D., Bloch, I., Sharon, E., Schueler-Furman, O., Wiener, R., and Tabach, Y. (2021). CladeOScope: functional interactions through the prism of clade-wise co-evolution. NAR Genom Bioinform 3, lqab024.

Wallis, N., Gershon, T., Shaaby, S., Oberman, F., Grunewald, M., Duran, D., Singh, A., Vainer, G., Spiegelman, V.S., Sharma, A.K., et al. (2025). AVJ16 inhibits lung carcinoma by targeting IGF2BP1. Oncogene 44, 3239–3254.

Wallis, N., Oberman, F., Shurrush, K., Germain, N., Greenwald, G., Gershon, T., Pearl, T., Abis, G., Singh, V., Singh, A., et al. (2022). Small molecule inhibitor of Igf2bp1 represses Kras and a pro-oncogenic phenotype in cancer cells. RNA Biol 19, 26–43.

Weidensdorfer, D., Stohr, N., Baude, A., Lederer, M., Kohn, M., Schierhorn, A., Buchmeier, S., Wahle, E., and Huttelmaier, S. (2009). Control of c-myc mRNA stability by IGF2BP1-associated cytoplasmic RNPs. RNA 15, 104–115.

Welshhans, K., and Bassell, G.J. (2011). Netrin-1-induced local beta-actin synthesis and growth cone guidance requires zipcode binding protein 1. J Neurosci 31, 9800–9813.

Winkler, F., Venkatesh, H.S., Amit, M., Batchelor, T., Demir, I.E., Deneen, B., Gutmann, D.H., Hervey-Jumper, S., Kuner, T., Mabbott, D., et al. (2023). Cancer neuroscience: State of the field, emerging directions. Cell 186, 1689–1707.

Xu, J., Wang, Y., Ren, L., Li, P., and Liu, P. (2025). IGF2BP1 promotes multiple myeloma with chromosome 1q gain via increasing CDC5L expression in an m(6)A-dependent manner. Genes Dis 12, 101214.

Yang, C.P., Samuels, T.J., Huang, Y., Yang, L., Ish-Horowicz, D., Davis, I., and Lee, T. (2017). Imp and Syp RNA-binding proteins govern decommissioning of Drosophila neural stem cells. Development 144, 3454–3464.

Yaniv, K., Fainsod, A., Kalcheim, C., and Yisraeli, J.K. (2003). The RNA-binding protein Vg1 RBP is required for cell migration during early neural development. Development 130, 5649–5661.

Yisraeli, J.K., Sokol, S., and Melton, D.A. (1990). A two-step model for the localization of a maternal mRNA in Xenopus oocytes: Involvement of microtubules and microfilaments in translocation and anchoring of Vg1 mRNA. Development 108, 289–298.

Zhang, J.Y., Chan, E.K., Peng, X.X., and Tan, E.M. (1999). A novel cytoplasmic protein with RNA-binding motifs is an autoantigen in human hepatocellular carcinoma. J Exp Med 189, 1101–1110.

Zhou, X., Li, G., Su, X., Wang, Y., Liu, L., Yang, C., Xu, Y., He, J., and Zhang, J. (2026). The role of m6A reader IGF2BPs in tumor metabolic reprogramming. Biochem Pharmacol 243, 117452.

